# Diversity in rest-activity patterns among Lake Malawi cichlid fishes suggests novel axis of habitat partitioning

**DOI:** 10.1101/2020.07.14.203505

**Authors:** Evan Lloyd, Brian Chhouk, Alex C. Keene, R. Craig Albertson

## Abstract

Animals display remarkable diversity in rest and activity patterns that are regulated by endogenous foraging strategies, social behaviors, and predator avoidance. Alteration in the circadian timing of activity or the duration of rest-wake cycles provide a central mechanism for animals to exploit novel niches. The diversity of the 3000+ cichlid species throughout the world provides a unique opportunity to examine variation in locomotor activity and rest. Lake Malawi alone is home to over 500 species of cichlids that display divergent behaviors and inhabit well-defined niches throughout the lake. These species are presumed to be diurnal, though this has never been tested systematically. Here, we measure locomotor activity across the circadian cycle in 12 cichlid species from divergent lineages and distinct habitats. We document surprising variability in the circadian time of locomotor activity and the duration of rest. In particular, we identify a single species, *Tropheops* sp. “red cheek” that is nocturnal. Nocturnal behavior was maintained when fish were provided shelter, but not under constant darkness, suggesting it results from acute response to light rather than an endogenous circadian rhythm. Finally, we show that nocturnality is associated with increased eye size, suggesting a link between visual processing and nighttime activity. Together, these findings identify diversity of locomotor behavior in Lake Malawi cichlids and provide a system for investigating the molecular and neural basis underlying the evolution of nocturnal activity.

## 1. Introduction

Animals display remarkable diversity in rest and activity patterns. Circadian differences in locomotor activity and rest can differ dramatically between closely related species, or even between individuals of the same species, raising the possibility that it can be adaptive and subject to selection[1–3]. Indeed, circadian regulation of locomotor activity is strongly associated with foraging strategies, social behaviors, and predator avoidance that are critical factors in organismal fitness[4,5]. Alteration in the circadian timing of activity or the duration of rest-wake cycles provide a central mechanism for animals to exploit novel niches.

Across phyla the timing of rest and activity is regulated by a circadian clock that persists under constant conditions, as well as acute response to environmental cues that include light and food availability[6]. For example, many teleost species display robust diurnal locomotor rhythms including the goldfish (*Carassius Auratus*), the Mexican tetra (*Astyanax mexicanus*), and the zebrafish (*Danio rerio*)[1,7,8]. Conversely, limited examples of nocturnal teleosts have been identified including the plainfin midshipman, the Senegalese sole, and the doctor fish, *Tinca tinca* [9–11]. Despite these conspicuous differences, variation in rest and activity patterns have not been well described within a lineage. Moreover, the ecological basis of such variation, and its relationship to niche exploitation has not been studied systematically.

Cichlids represent a leading model for investigating the evolution of development, morphology, and complex behavior. In Lake Malawi alone, there are over 500 species of cichlids, inhabiting a diversity of environmental and feeding niches. Cichlid species exhibit a high degree of habitat fidelity and partition their environment along discrete ecological axes, including distinct biotic (food availability, predation, and parasites) and abiotic (light, water chemistry) environments that play a critical role in the origins and maintenance of cichlid biodiversity[12–17]. Predation on Malawi cichlids is considered to be relatively low, which is thought to have contributed to their evolutionary and ecological success[18]. However, the lake is home to predators, including the Cornish jack *Mormyrops anguilloides*, that feed on cichlids in the intermediate and near-shore rocky habitat. *M. anguilloides* are weakly electric fish that hunt at night using electrical pulses thought to be undetectable by cichlids[19]. Field studies on this predatory behavior have suggested that cichlids are largely diurnal[19], in agreement with the notion that rest represents a form of adaptive inactivity that allows for predator avoidance[20]. Deviations from diurnal activity have been noted for new world cichlids, which exhibit nocturnal parental care of eggs[21,22], and the ability of some Malawi cichlids to forage in low-light conditions, via widened lateral line canals, suggests the potential for nocturnal behaviors to evolve in this group[23]. Given that Malawi cichlids exhibit an impressive magnitude of diversity in an array of anatomical and behavioral traits, we reasoned that they may also exhibit variation in rest-activity patterns. Indeed, this could represent an important, but underappreciated, dimension of habitat partitioning.

The development of automated tracking of locomotor activity in fish species has been applied for the study of sleep and locomotor activity in zebrafish and Mexican cavefish[24]. These methodologies provide the opportunity for comparative approaches that examine differences in activity between populations, and across contexts. Here, we extend this methodology to study sleep across 12 species of cichlids, from diverse habitats. Our choice of species focuses on the near-shore rock-dwelling clade of Malawi cichlids (i.e., *mbuna*), but we also include representative species from other major lineages. Our goal is not to characterize the evolution of rest-activity patterns *per se*, but rather to better understand the degree and type of variation exhibited by this group. We identify robust variation in the quantity, as well as the circadian timing, of rest and activity. In addition, this analysis reveals, for the first time, a nocturnal species of Malawi cichlid, suggesting that circadian regulation of activity may provide a mechanism for niche exploitation in African cichlids. Together, these findings suggest cichlids can be used as a model to study the evolution of molecular mechanisms for variation in locomotor rhythms.

## 2. Materials and Methods

### (a) Fish stocks and husbandry

Cichlids used for experiments were reared following standard protocols approved by the University of Massachusetts Institutional Animal Care and Use Committee. Cichlids were housed in the Albertson fish facilities at a water temperature of 28.5°C, kept on a 14:10 hour light-dark cycle, and fed a diet of a flake mixture consisting of ~75% spirulina algae flake and ~25% yolk flake twice a day. Cichlids were derived from wild-caught animals that were either reared in the Albertson fish facilities (*Labeotropheus trewavasae, Maylandia zebra, Tropheops sp.* “red cheek”), or obtained through the aquarium trade (*Sciaenochromis fryeri, Copadichromis trewavasae, Aulonocara stuartgranti, Dimidiochromis compressiceps, Labeotropheus fuelleborni, lodotropheus sprengerae, Tropheops* sp. “red fin”, *Tropheops* sp.“elongatus Boadzulu”). Because of the nature of the testing tanks (see below), all fish were tested at the late juvenile stage, making sex determination difficult to assess at the time; however, after the experiments took place, stocks were grown out and it could be confirmed that sex ratios were 50:50 on average.

### (b) Behavioral analysis

24 hours prior to the beginning of each experiment, juvenile fish were transferred from their home tanks into 10L tanks (Carolina Biologicals) with custom-designed partitions that allowed for up to 3 fish to be individually housed in each tank. After 24 hours of acclimation, fish were fed, tanks were given a 50% water change to maintain water quality, and behavior was recorded for a 24 hour period beginning at zeitgeber time (ZT) 3, 3 hours after light onset. Videos were recorded at 15 frames/second using a USB webcam (LifeCam Studio 1080p HD Webcam, Microsoft) through the video processing software VirtualDub (v1.10.4). To allow for recording during the dark period and provide consistent lighting throughout the day, cameras were modified by removing their IR filters and replacing with IR long-pass filters (Edmund Optics Worldwide), and tanks were illuminated from behind using IR light strips (Infrared 850 nm 5050 LED Strip Light, Environmental Lights).

For experiments testing the effect of shelter on locomotor activity, a small PVC tube, 3” in length x 1” outer diameter, was added to each chamber at the beginning of the acclimation period. For experiments testing the effect of light, fish were acclimated to their tanks on a normal 14:10 LD cycle, and then recorded in 24 hours of darkness.

Following acquisition, recordings were processed in Ethovision XT 15 (Noldus) to extract positional data for individual fish throughout the 24 hour period, and this data was used to calculate velocity and locomotor activity, as previously described[25].

### (c) Analysis of activity and rest

To identify variation in rest and activity patterns across cichlid species, positional data was exported from Ethovision and analyzed using a custom-made Perl script (v5.10.0) and Excel Macro (Microsoft). A threshold of 4 cm/sec was set to correct for passive drift of the animal; any reading over this threshold was classified as active swimming and used to calculate velocity. Any period of inactivity lasting greater than 60 seconds was classified as a “rest” bout, and the time and duration of each rest bout was recorded to generate profiles of rest throughout the day.

### (d) Measurements of eye size

Fish were imaged using a digital camera (Olympus E520) mounted to a camera stand. All images included a ruler. Using the program Image J[26], measures of standard length, head length, and eye area were obtained for each fish. Eye size was measured in fish used in the behavioral analysis. In addition, when possible, we augmented these samples with wild-caught animals from the Albertson lab collections. In particular, we added wild-caught samples to the *L. fuelleborni, M. zebra, T. sp.*“red cheek”, and *T. sp.* “red fin” populations. Two measures of eye-size were obtained: total area, and area relative to head length. Because relative eye-size exhibits strong allometric effects[27],residuals were obtained via a linear regression of eye-size on standard length in R[28]. All statistical analyses were based on residuals. Results were the same whether we used absolute eye-size or eye-size relative to head length. We therefore only present data using absolute eye-size.

### (e) Statistics and analysis

One-Way ANOVAs were carried to identify inter-species differences in overall locomotor activity, average waking velocity, rest duration, and total time in shelter between species. To identify differences between multiple conditions, such as activity in the light vs. dark, or shelter vs no-shelter conditions, a Two-Way ANOVA was carried out, and followed by Sidak’s multiple comparisons *post-hoc* test. To identify significant rhythms in activity across the day-night cycle, an “activity change ratio” was calculated as follows: First, average hourly day- and night-time activity were calculated for each fish. Night-time activity was subtracted from day-time activity, and the result was divided by their sum, providing a normalized day/night preference score. To identify significant rhythmicity, one-sample t-tests were performed. To identify differences between the *mbuna* and *non-mbuna* groups, nested ANOVAs were performed. All statistical analyses were carried out using InStat software (GraphPad Prism 8).

## 3. Results

### (a) Variation in activity and rest behaviors

To measure variation in activity across Lake Malawi cichlids, we compared the locomotor activity in twelve different species, across eight genera, of cichlids that were selected for diversity in habitat, behavior, and lineage representation. We sampled more deeply in the rock-frequenting *mbuna* clade (n=7 species; n=4 genera), which occupy a complex, three-dimensional habitat characterized by a high density of cichlid individuals (Fig 1A). In addition, we analyzed activity patterns in four *non-mbuna* species, which occupy the intermediate to open-water habitat (Fig 1B). Following an initial 24 hour period of acclimation, activity was recorded in individually housed juvenile fish across 24 hours in standard light-dark conditions, with infrared lighting used to monitor locomotor activity during the night as previously described in *A. mexicanus*[25]. Quantification of total locomotor activity over 24 hours identified marked variation across species, with certain species (i.e., *S. fryeri*) exhibiting significantly lower activity than all other species tested, while the activity of others (i.e., *T.* sp. “elongatus Boadzulu”) was significantly greater than all other species (Fig 1C). Notably, variation in mean activity was continuously distributed between these two extremes. In addition, there was a division between *mbuna* and non-*mbuna* species, with *mbuna* species trending towards increased locomotion relative to *non-mbuna* species (*p* = 0.052).

**Fig 1.**
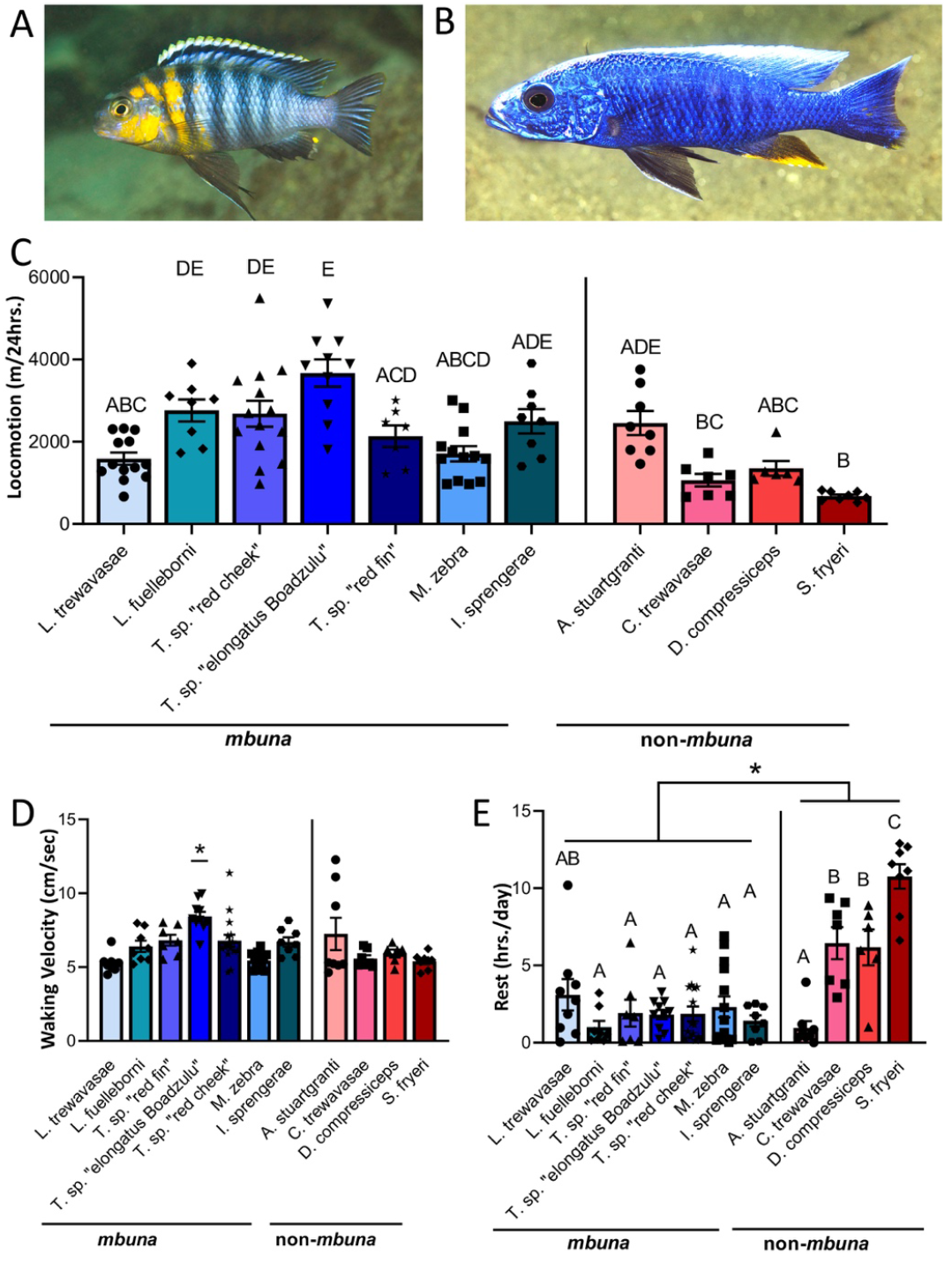
Evolved differences in locomotor activity between cichlid species. (A) Morphology of *T.* sp. “red cheek” of the *mbuna* group. (B) Morphology of *S. fryeri* of the *non-mbuna* group. Images by Ad Konings, Cichlid Press. (C) Total locomotor activity over 24 hours varies significantly across 11 cichlid species (one-way ANOVA: *F*_10, 91_ = 11.10, *p*<0.0001). *Mbuna* species trend towards higher activity than *non-mbuna* species, although this relationship does not reach significance (nested ANOVA, *F*_1,9_ = 5.009). (D) Waking velocity over 24 hours is significantly elevated in only one species of cichlid, *Tropheops* sp. “elongatus Boadzulu” (one-way ANOVA, F_10,89_ = 5.398). (E) Consolidated periods of rest (>60 seconds) vary significantly across *mbuna* and non-*mbuna* groups. (nested ANOVA, *F*_1,9_ = 7.808, *p* = 0.0209).

To determine whether these differences were due to hyperactivity or differences in rest, we measured the average waking velocity for each population. Among all species tested, only one (*T.* sp. “elongatus Boadzulu”), displayed significantly higher swimming velocity, suggesting the bulk of the variation among species is due to differences in rest/activity regulation (Fig 1D). In agreement with this notion, there were significant inter-species differences in the duration of rest bouts lasting greater than one minute (Fig 1E). The majority of species displayed very little rest, averaging less than 3 hours/day, while three species, *C. trewavasae, D. compressiceps*, and *S. fryeri* (all *non-mbuna*) spent significantly longer resting than all other species tested. The average rest duration of *S. fryeri* was over 10-fold different than other species tested. Together, these findings suggest that differences in total locomotor activity between cichlid species are largely attributable to differences in rest. Notably, *mbuna* species together rested significantly less than non-*mbuna* species (Fig 1E), possibly reflecting adaptation to the near-shore rocky habitat. Support for this possibility, as opposed to historical contingency, is the observation that *A. stuartgranti*, a *non-mbuna* species that co-occurs with *mbuna*, rests less than other non-*mbuna* species (Fig 1E).

### (b) Variation in patterns and magnitudes of rhythmic activity

To determine whether there are differences in circadian modulation of activity, we compared activity over the light-dark cycle (Fig 2A). We found evidence for strong diurnal activity in three *mbuna* species (*L. fuelleborni*, *T*. sp. “elongatus Boadzulu”, and *l. sprengerae*), while activity did not significantly differ based on light or dark phases in seven species tested (Fig 2B). A single species, *Tropheops* sp*. “*red cheek*”*, was significantly more active in the night, providing the first evidence for nocturnality in a Lake Malawi cichlid (Fig 2B). To account for variation in total locomotion between fish of different species, we quantified preference for light and dark activity for each individual tested. In agreement with quantification of average locomotor activity, *T.* sp. *“*red cheek*”*had significantly greater preference for nighttime activity while *L. fuelleborni, T.* sp. “elongatus Boadzulu”, and *I. sprengerae*, had significantly greater preference for daytime activity (Fig 2C). This analysis also suggests a preference for diurnal activity in *L. trewavasae*, and for nocturnal activity in two additional *non-mbuna* species (*C. trewavasae* and *S. fryeri*).

**Fig 2.**
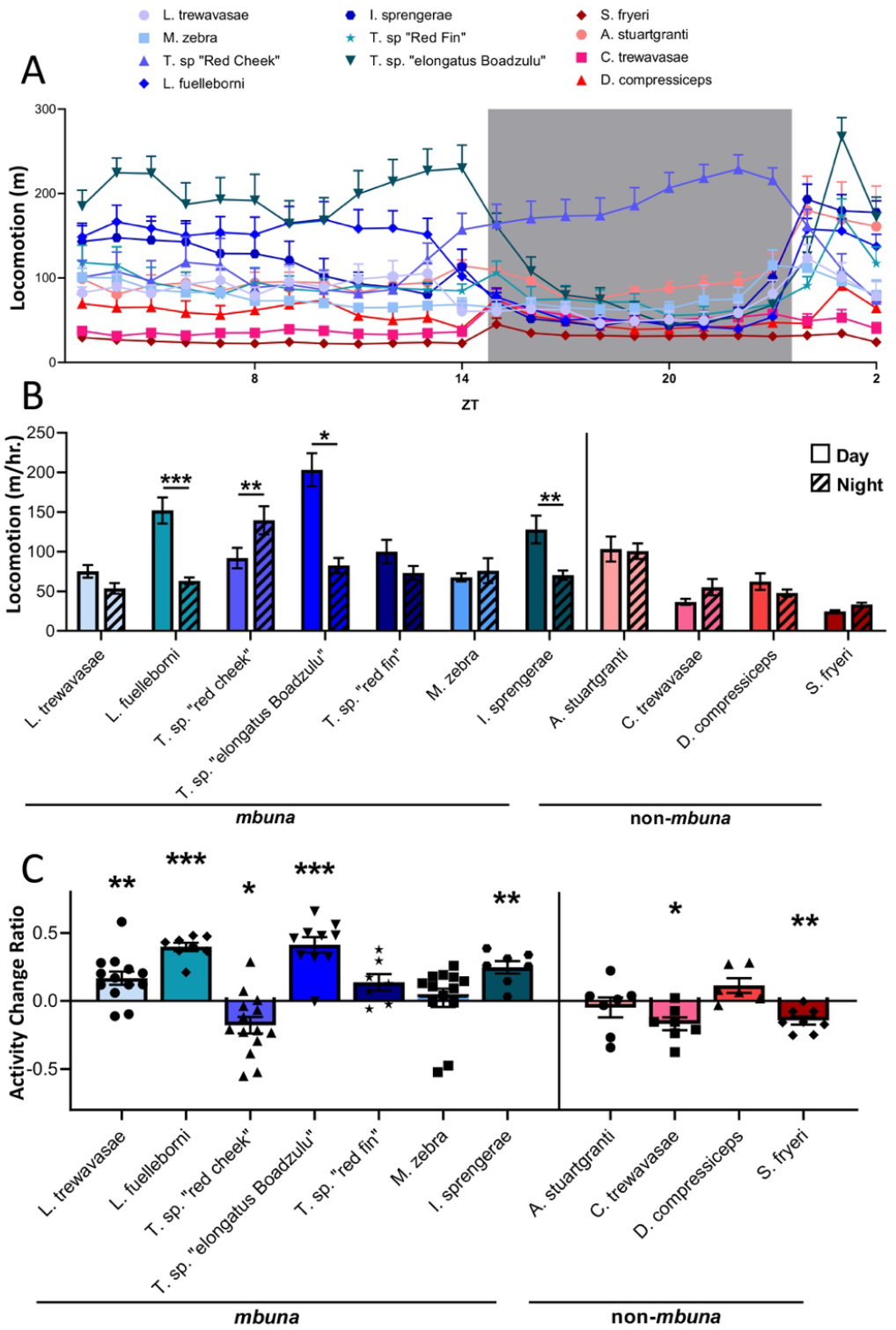
Variation in daily activity rhythms across the day/night cycle. (A) Activity profiles of all species tested across 24 hours, beginning at ZT3. (B) Several species exhibit significantly increased activity during the subjective day (*L. fuelleborni, T.* sp. “elongatus Boadzulu”, *I. sprengerae*) while a single species exhibits increased locomotor activity during the subjective night (*T.* sp. “red cheek”) (two-way ANOVA, *F*_10,91_ = 13.27). (C) Activity change scores, calculated as the difference between daytime and nighttime activity, divided by their sum, reveal differences in day/night preference across cichlid species (one-sample t-test; *L. trewavasae (t_12_ = 3.419, p* = 0.0051), *L. fuelleborni* (*t_7_* = 12.93, *p* < 0.0001), *T.* sp. “red cheek”(*t_13_* = 2.930, *p* < 0.0117), *T*. sp. “elongatus Boadzulu”(*t_9_* = 7.182, *p* < 0.0001), *I. sprengerae* (*t_6_* = 5.374, *p* = 0.0017), *C. trewavasae* (*t_6_ = 3.555*, p= 0.012), *S. fryeri* (*t_7_* = 4.693, *p* = 0.0022).

**I**n other diurnal teleosts, such as *A. mexicanus* and *D. rerio*, rest is largely consolidated during nighttime. To quantify time of day differences in rest across cichlid species, we compared the average amount of rest per hour across the 14 hours of light and 10 hour dark periods (Fig S1). This analysis is largely in agreement with analysis of locomotor activity, with day-active species consolidating rest during the dark period, and *vice versa*.

Since *T.* sp. “red cheek” is a highly territorial and aggressive species[29,30], it is possible that the nighttime activity of this species represents a search strategy for locations that provide shelter from predators, as opposed to a natural reflection of activity patterns. To differentiate between these possibilities, we provided each animal with a 3 inch cylindrical shelter (PVC piping), and measured behavior across light and dark conditions (Fig 3A). We analyzed the total activity across the circadian cycle, as well as time spent in the shelter in *T.* sp. “red cheek”, as well as in *L. trewavasae* and *M. zebra*, closely related *mbuna* species that co-occur with *T.* sp. *“*red cheek*”*. These two species also exhibited lower and indistinguishable activity levels during the day and night, and we were interested to see if the addition of shelter would alter this pattern. When provided a hiding spot, *T.* sp. “red cheek” remained robustly nocturnal, while *M. zebra* and *L. trewavasae* did not show light dark preference, which is consistent with their activity patterns in the absence of shelter (Fig 3B). We quantified the total time animals spent within the shelter and found that *L. trewavasae* spent significantly more time in the shelter than *M. zebra* and *T.* sp. “red cheek” (Fig 3C), which is consistent with this species’ behavior in the wild. *L. trewavasae* has an elongated and dorsal-ventrally compressed body plan, and exhibits habitat preference for cracks and crevices in the wild[30,31]. Further, *L. trewavasae* spent more time in the shelter during the night period, consistent with an increased need to avoid nocturnal predators (Fig 3D). Conversely, there were no differences in shelter preference between light or dark periods for *M. zebra* and *T.* sp. *“*red cheek”. Together, these findings suggest that the presence of a shelter does not significantly impact the activity pattern of the cichlid species tested, and that the nocturnal locomotor activity of *T.* sp. “red cheek” does not represent a search for shelter.

**Fig 3.**
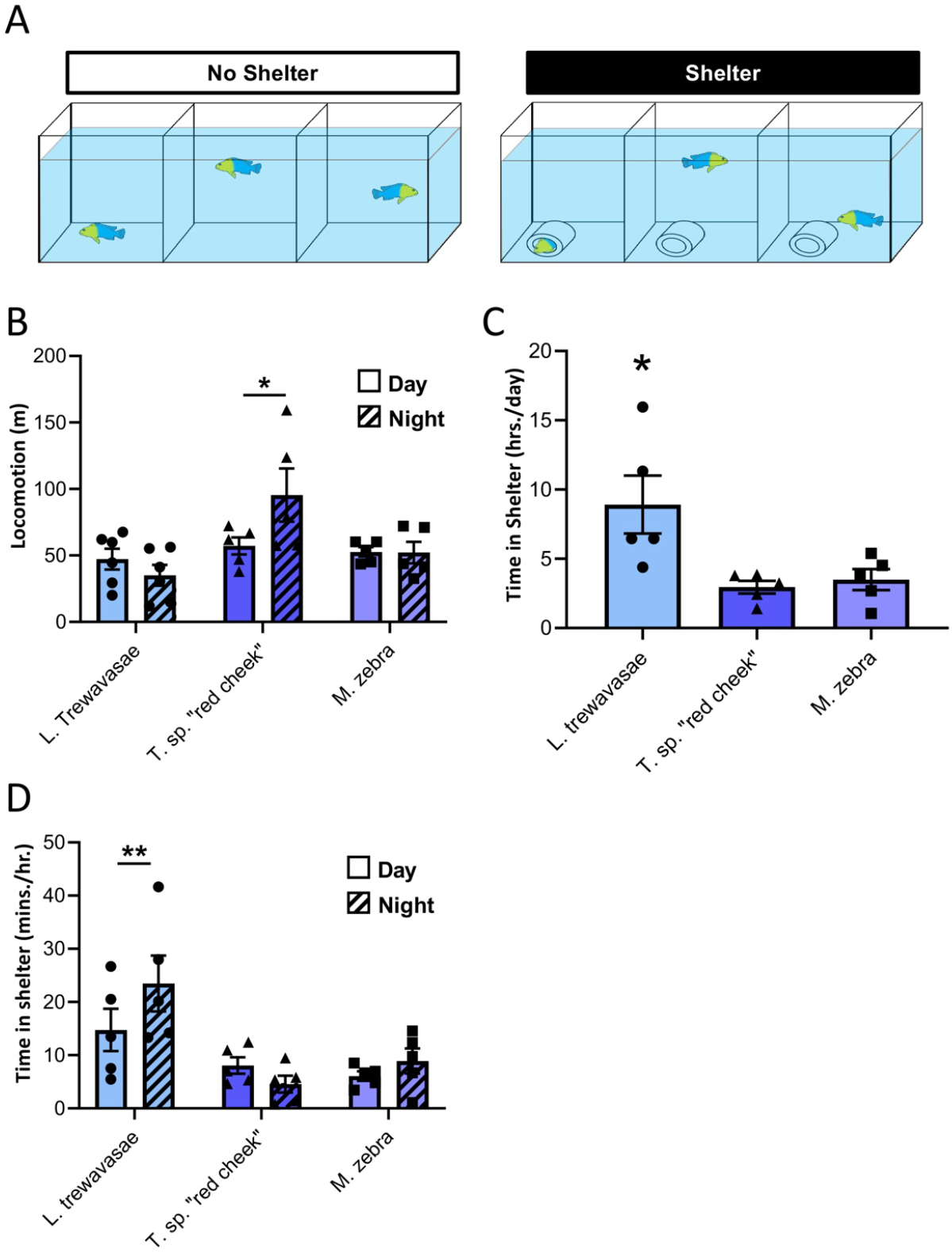
Presence or absence of shelter does not affect the nocturnal phenotype of *T.* sp. “red cheek”. (A) Select cichlid species were tested for locomotor activity over 24 hours in the presence or absence of a 3 inch PVC tube, providing the option to take shelter at any point during the day. (B) *T.* sp. “red cheek” maintain bias for nocturnal activity in the presence of shelter (two-way ANOVA, *F*_2,12_ = 7.9, *p* = 0.0065). (C) *L. trewavasae* exhibit significantly greater preference for the shelter relative to other species tested, consistent with knowledge of the species’ ecological niche (one-way ANOVA, *F*_2,12_ = 6.305, *p* = 0.0134). (D) Preference for shelter increases at night only in *L. trewavasae* (two-way ANOVA, *F*_2,12_ = 7.9, *p* = 0.0065).

It is possible that the nocturnal locomotor behavior of *T.* sp. “red cheek” is due to an endogenous circadian rhythm or a differential response to light. To distinguish between these possibilities, we measured locomotor activity under constant dark conditions. Briefly, fish were acclimated under standard 14:10 light dark conditions, then activity was recorded for 24 hours under constant darkness (Fig 4A). While *T.* sp. “red cheek” are significantly more active during the dark period under light:dark conditions, there was no difference between light and dark activity under constant darkness. (Fig 4A). A comparison of total activity between the day (with light present) and the subjective day (darkness) reveals that activity is significantly lower in the presence of light (Fig 4B). These findings are consistent with a role for light in suppressing activity, thereby inducing nocturnal behavior.

**Fig 4.**
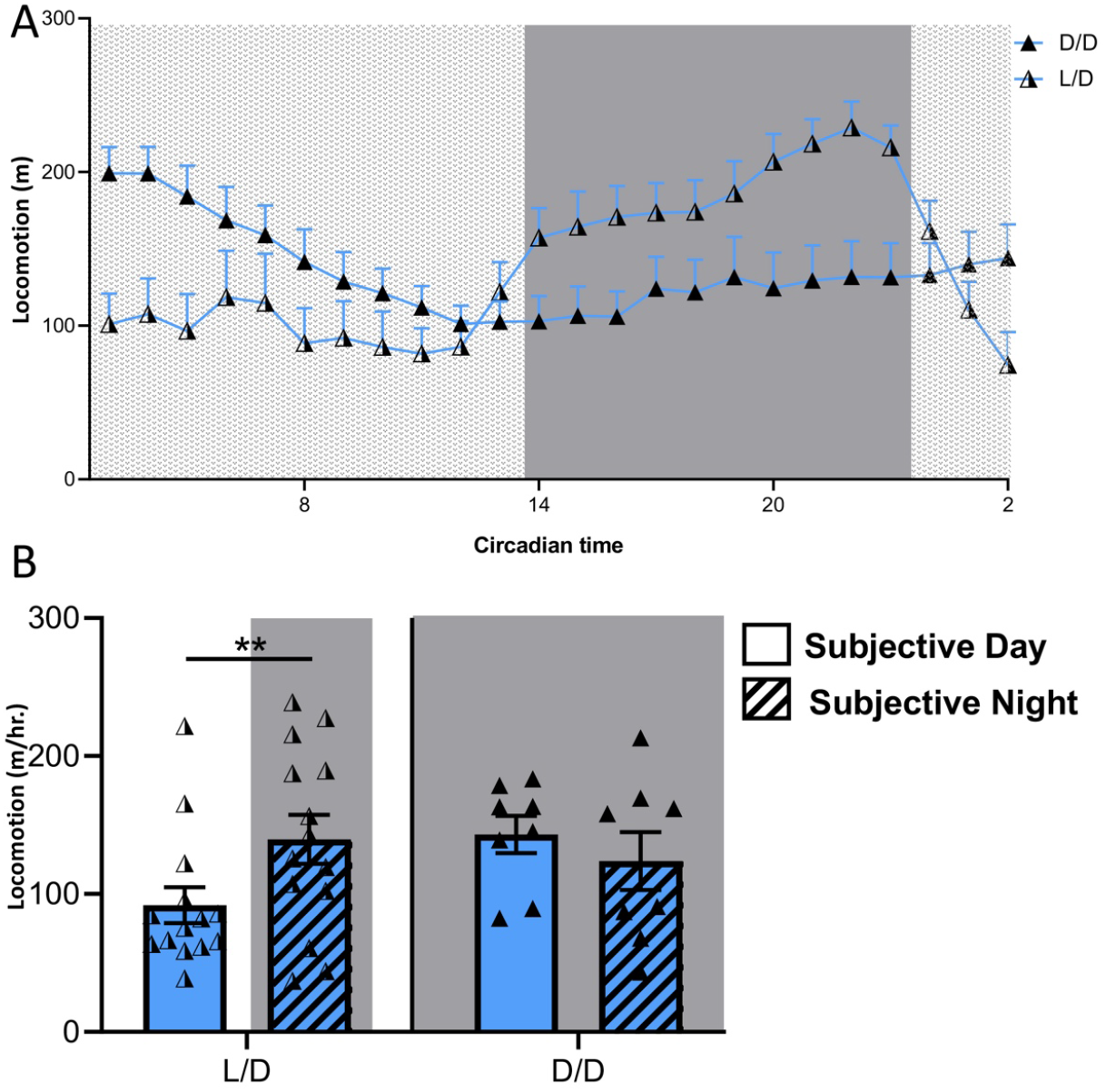
Locomotor activity in *T.* sp. “red cheek” is modulated by the presence/absence of light. (A) Activity profiles of *T.* sp. “red cheek” under a 14:10 LD cycle (half-shaded triangles), and in constant darkness (fully shaded triangles). (B) Under a 14:10 LD cycle, *T.* sp. “red cheek” increase activity during the day; in 24 hours of darkness, activity remains elevated throughout the 24-hour period (two-way ANOVA, *F*_1,20_ = 7.393, *p* = 0.0132).

### (c) Activity is higher in territorial species

General information regarding each species’, habitat, behavior, prey-preference, and phylogenetic relationship is provided in Table 1. To determine whether any variables of rest or activity associate with these ecological factors, we compared locomotor data with known ecological variables. Unsurprisingly, species described as territorial exhibited, on average, greater overall activity compared to those characterized as weakly or non-territorial. We note, however, that any conclusion about the relationship between locomotor activity and ecology/phylogeny may be premature, as significant differences in rest-activity behavior exist between closely related and ecologically similar species (e.g., within *Tropheops* and *Labeotropheops*). The more general conclusion to be drawn from these data is that Lake Malawi cichlids exhibit substantial and continuous variation in activity levels and patterns.

**Table 1.** General information about the Lake Malawi species under study. Information based on [30,31]. Abbreviations: G = generalist, S = specialist, N = non-territorial, T = territorial, W = weakly territorial, SL = standard length.

| Genus | species | Clade | Habitat | Depth distribution | Prey Type | Specialist/<br>Generalist |  | Territorial | Size (SL) | Distribution |
| --- | --- | --- | --- | --- | --- | --- | --- | --- | --- | --- |
| *Aulonocara* | *stuartgranti* | Deep Benthic | Intermediate | 5-15m | Benthic invertebrates | S |  | T | <10cm | Lake-wide |
| *Copadichromis* | *trewavasae* | Utaka | Intermediate/open water | 10-30m | Zooplankton | S |  | T | <10cm | Lake-wide |
| *Dimidiochromis* | *compressiceps* | Shallow Benthic | Intermediate with plants | <30m | Small fish/fry | S |  | W | 10-20cm | Lake-wide |
| *Sciaenochromis* | *fryeri* | "Old" sand dweller | Intermediate | 10-40m | Small fish/fry | S |  | W | 12-14cm | Lake-wide |
| *Tropheops* | sp. "red fin" | *mbuna* | Rocks with sediment | >10m | Algae/detritus | G |  | T | <10cm | Lake-wide |
| *Tropheops* | sp. "elongatus<br>Boadzulu" | *mbuna* | Rocks with sediment | >10m | Algae/plankton/<br>detritus | G |  | T | <10cm | Specific locations,<br>south |
| *Tropheops* | sp. "red cheek" | *mbuna* | Sediment-free rocks | <10m | Attached filamentous<br>algae | S |  | T | <10cm | Specific locations,<br>south and north<sup>2</sup> |
| *Labeotropheus* | *trewavasae* | *mbuna* | Between/under rocks<br>with sediment | 0-20m | Algae/detritus | G |  | W | <10cm | Lake-wide |
| *Labeotropheus* | *fuelleborni* | *mbuna* | Sediment-free rocks | <10m<sup>1</sup> | Attached filamentous<br>algae | S |  | T | 10+cm | Lake-wide |
| *Maylandia* | *zebra* | *mbuna* | Diverse rocky habitat | 5-20m | Loose algae | G |  | T | <10cm | Lake-wide |
| *Iodotropheus* | *sprengerae* | *mbuna* | Diverse rocky<br>and intermediate | 3-15m<sup>1</sup> | Diverse omnivorous | G |  | N | <10cm | Southeast |

<sup>1</sup> These species have been observed to penetrate much deeper waters (e.g., ~40m).
<sup>2</sup> This distribution pattern suggests a lake-wide historical distribution.

### (d) Eye size is associated with night-time activity

Across fish species, nocturnality or adaptation to low-light conditions is associated with larger eye size. In addition, species that rely on visual modes of foraging generally develop larger eyes[32–34]. On the other hand, species adapted to forage on attached algae generally possess smaller eyes, consistent with a functional tradeoff for the production of power during jaw closure[35]. Specifically, algal scrapers tend to exhibit smaller and dorsally shifted eyes to accommodate larger adductor muscles that are situated below the eyes[36]. To understand how eye size relates to these variables, we measured eye size in cichlid individuals in all species tested (Fig 5A, 5B), and tested for significant correlations. Notably, we did not observe an obvious association between eye size and lineage or foraging mode (Fig 5B). While the visual hunting species *C. trewavasae* and *S. fryeri* possess larger eyes on average, *D. compressiceps*, an ambush-hunter, has the smallest eyes of the species measured. Likewise, while the algal scraping species within the genus *Labeotropheus* has relatively small eyes, the attached algae specialists, *T.* sp. “red cheek”, has the largest relative eye size of the species measured. The other species with large eyes was *A. stuartgranti*, which is a sonar hunter with enlarged lateral line canals capable of foraging in low-light conditions[23]. Neither did we identify a correlation between rest amount and eye size (Fig 5C). However, there was a strong correlation between eye size and preference for night-time activity (Fig 5D). Whether the large eye size in these species represents an adaptation to nocturnality remains to be tested, but it is a notable morphological correlate worthy of further investigation.

**Fig 5.**
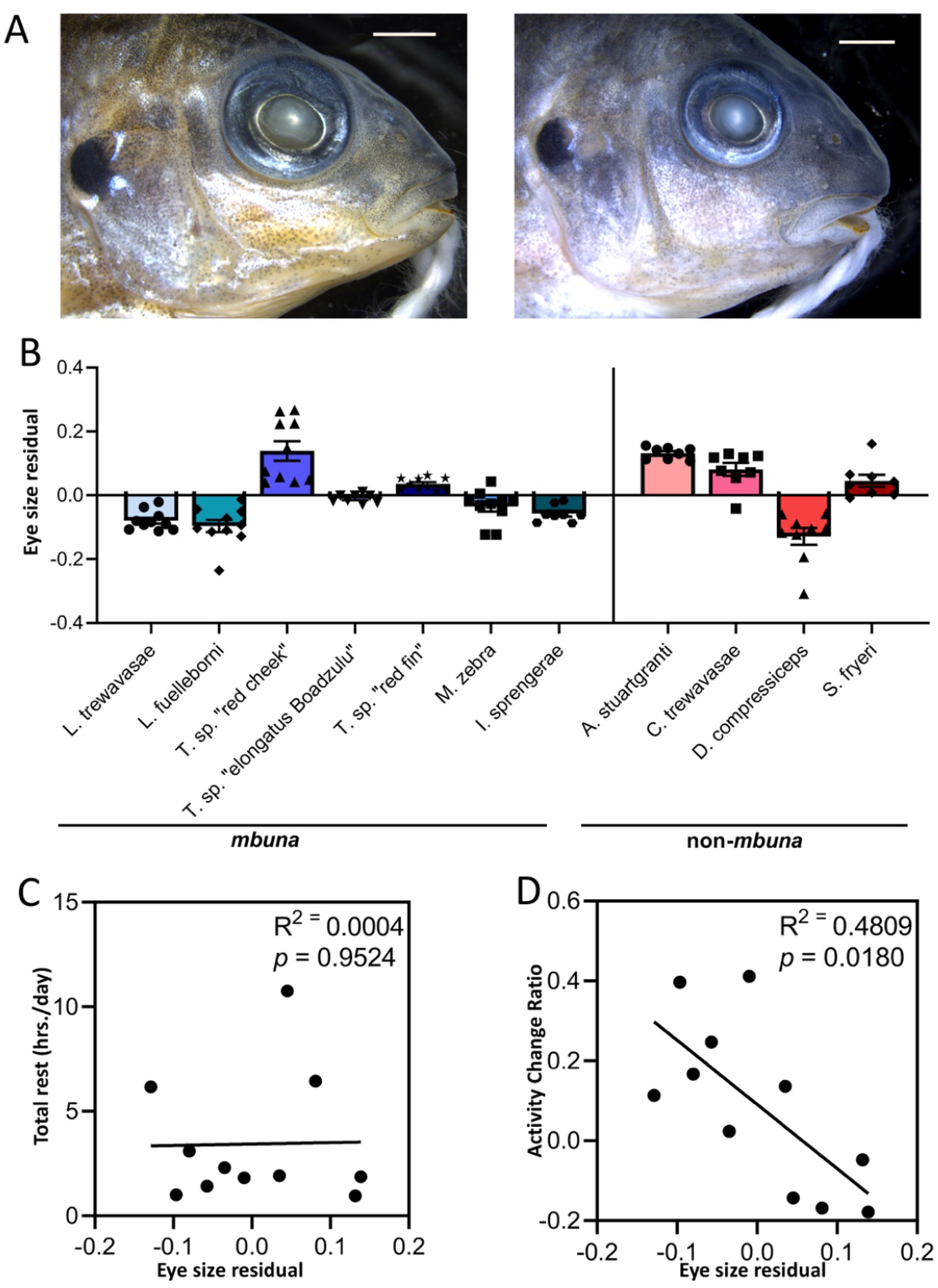
(A) Photographs demonstrating difference in relative eye size between *T.* sp. “red cheek”, left, and *L. fuelleborni*, right. Both specimens were approximately 4 months old. Scale bars equal 2mm. (B) Variation in eye size across species. (C) There is no relationship between eye size and total rest amount. (D) There is a significant negative relationship between eye size and a bias towards daytime activity.

## 4. Discussion

The diversity of the ~3000 cichlid species throughout the world provides a unique opportunity to examine the effects of ecological niche and evolutionary history on the regulation of locomotor activity and rest. Cichlid species have undergone adaptive radiations, resulting in morphologies and behaviors that can be highly specialized to specific ecological niches. The ecology inhabited by cichlids includes species with habitat fidelity to shorelines, deep water, and the intermediate zone between rocky and sandy regions. In addition, many species are generalists and inhabit multiple different niches. Here we focused our analysis on Lake Malawi cichlids, that alone likely contains over 500 species of fish, many of which share overlapping ecological niches[37]. The well characterized ecosystem within the lake, as well as the taxonomic diversity uniquely positions cichlids for investigating the role of ecology in shaping behavioral evolution. Indeed, an important outstanding question is how can so many species with dietary overlap co-exist in this lake? Many factors have been proposed to contribute including the multitude of ecological resources available in this large tropical lake, low predation, as well as the ability of cichlid species to evolve highly specific courtship and feeding behaviors[18,38]. Circadian regulation of activity and rest may provide an additional contributor to niche partitioning, reproductive isolation and even speciation, yet these behaviors have not been studied systematically. The finding that the timing and duration of rest and activity varies dramatically, and continuously, between populations of Lake Malawi cichlids suggest this is a fruitful line of inquiry.

While circadian rhythms have been studied in detail across many different animal species, surprisingly little is known about the presence and regulation of free running rhythms in teleosts. For example, Nile tilapia, *O. niloticus*, display extreme variability under light-dark conditions that ranges from diurnal to nocturnal, yet the majority of animals maintain rhythms of ~24hrs under constant dark conditions[39]. Feeding is likely a critical mediator of activity rhythms, though in some species, the daily timing of feeding differs from locomotor activity. For example, zebrafish are highly diurnal and maintain 24 hour rhythms, yet feeding occurs primarily during the night[40]. A similar trend has been noted in cichlids, where diurnal species exhibit mating and brooding behaviors primarily at night[21,41]. These findings suggest a high degree of flexibility in the circadian regulation of behavior, and that the circadian timing of many behaviors may differ from locomotor behavior that is typically used as a primary readout of the circadian clock[42]. Here, we focused specifically on locomotor activity and did not provide social conspecifics or food that could influence the timing of activity. Fully understanding the evolution circadian behavior of each species and its relationship to its natural environment will require examining additional behaviors that may be under circadian regulation.

A notable finding from this study is a species that appears to be nocturnal. *Tropheops sp.*“red cheek” is a member of a highly speciose and ecologically diverse lineage[13,30,43]. It is a vigorously territorial species that occupies the near shore rocky habitat, where males defend large patches of rocks, cultivating algae gardens that they only allow potential mates to feed from. This species exhibits significant habitat and dietary overlap with *L. fuelleborni*, another algae foraging species from the rocky shallows. *L. fuelleborni* is arguably one of the most ecologically successful species in the lake, with numerous anatomical adaptations that enable it to dominate this niche[31,44–46]. How then might another species coexist with such a well-adapted forager? Based on the results presented here, it is tempting to speculate that *L. fuelleborni* and *T. sp.*“red cheek” are partitioning their habitat by rest-activity patterns. Consistently, these two species (1) are among the most active of any measured, (2) are both strongly rhythmic, and (3) their rhythmicity is opposite of one another.

Our findings raise the possibility that *T. sp. “* red cheek” is nocturnal in the wild, and the limited amount of night filming that has been performed in Lake Malawi supports this notion. Specifically, Arnegard and Carlson (2005) documented the nocturnal predatory behavior of the weakly electric species, *M. anguilloides*, on cichlids in the rocky habitat. The footage (available at https://malawicichlids.com/mw19000.htm), is impressive and shows the success of the “pack” hunting strategy employed by *M. anguilloides.* Two cichlid species (based on male breeding color) are readily apparent in the footage, *A. stuartgranti* and *T.* sp. “red cheek”. Indeed, the very first fish seen in the night footage is a male *T.* sp. “red cheek” (@ 1:15). This fish is not resting within a rocky cave, crack or crevice, but rather it is actively swimming well above the rocks. In fact, in the ~6 minutes of night footage, no fewer than 5 *T.* sp. “red cheek” individuals can be observed, many *A. stuartgranti* are observed as well. As a point of comparison, no *Labeotropheus* or *Maylandia* species are readily observed at night, though they are common in the day footage at the beginning and end of the ~8 minute film. This filming was not intended to address questions related to rest-activity patterns in cichlids, and so we are cautious about drawing firm conclusions; however, the trends are conspicuously consistent with our laboratory results.

It is important to note that our analyses are limited to rest, and we did not examine sleep *per se*. Across phyla, ranging from jellyfish to humans, sleep can be defined by shared behavioral characteristics that include consolidated periods of behavioral quiescence, homeostasis following deprivation and increased arousal threshold, and species-specific posture[47]. In teleosts, the duration of inactivity associated with sleep has been defined as one minute of immobility in larval *A. mexicanus* and zebrafish, and the same duration for adult *A. mexicanus[48,49].* The duration of sleep and rest is highly variable across many other teleost species, and even between individuals of the same species. For example, different populations of *A. mexicanus* display extreme differences in sleep and activity, with cave dwelling populations of *A. mexicanus* sleeping less than river-dwelling surface fish counterparts. These differences presumably evolved, at least in part, due to increased foraging needs in a nutrient-poor cave environment[50]. Based on previous work in fishes, we defined rest as the total duration of inactivity bouts longer than one minute, and therefore these phenotypes may reflect differences in sleep duration across cichlid species. While specifically examining sleep in cichlids will require defining the period of immobility associated with changes in arousal threshold, posture, poster, and other behavioral characteristics of sleep, we submit that it represents a fruitful line of inquiry as it offers an ideal system in which to delve further into the evolution of sleep and its molecular underpinnings.

## Data accessibility

Will be determined upon acceptance.

## Author’s contributions

A.C.K and R.C.A conceived the project. E.L. and B.C. ran the sleep experiments. R.C.A measured eye sizes. E.L. analyzed the behavioral and morphological data. A.C.K, E.L. and R.C.A interpreted the results and wrote the manuscript. All authors were involved with rounds of editing and proofing of the figures and manuscript text.

## Acknowledgements

We thank members of the Albertson and Keene laboratories for logistical support with fish care, as well as for thoughtful discussions on various aspects of this project. Ad Konings, Cichlid Press, provided the images of cichlid species in Figure 1. He is acknowledged for his continued and generous support of the cichlid research community through the use of his extensive image library.

## References

1. Duboué ER, Keene AC, Borowsky RL. 2011 Evolutionary Convergence on Sleep Loss in Cavefish Populations. Curr. Biol. 21, 671–676. (doi:10.1016/J.CUB.2011.03.020)

2. Brown EB, Torres J, Bennick RA, Rozzo V, Kerbs A, DiAngelo JR, Keene AC. 2018 Variation in sleep and metabolic function is associated with latitude and average temperature in *Drosophila melanogaster*. Ecol. Evol. 8, 4084–4097. (doi:10.1002/ece3.3963)

3. Hammond TT, Palme R, Lacey EA. 2018 Ecological specialization, variability in activity patterns and response to environmental change. Biol. Lett. 14, 20180115. (doi:10.1098/rsbl.2018.0115)

4. Vaze KM, Sharma VK. 2013 On the Adaptive Significance of Circadian Clocks for Their Owners. Chronobiol. Int. 30, 413–433. (doi:10.3109/07420528.2012.754457)

5. Siegel JM. 2005 Clues to the functions of mammalian sleep. Nature 437, 1264–1271. (doi:10.1038/nature04285)

6. Huang W, Ramsey KM, Marcheva B, Bass J. 2011 Circadian rhythms, sleep, and metabolism. J. Clin. Invest. 121, 2133–2141. (doi:10.1172/JCI46043)

7. Iigo M, Tabata M. 1996 Circadian rhythms of locomotor activity in the goldfish Carassius auratus. Physiol. Behav. 60, 775–781. (doi:10.1016/0031-9384(96)00131-x)

8. Zhdanova I V., Wang SY, Leclair OU, Danilova NP. 2001 Melatonin promotes sleep-like state in zebrafish. Brain Res. 903, 263–268. (doi:10.1016/S0006-8993(01)02444-1)

9. Feng NY, Bass AH. 2016 “Singing” Fish Rely on Circadian Rhythm and Melatonin for the Timing of Nocturnal Courtship Vocalization. Curr. Biol. 26, 2681–2689. (doi:10.1016/j.cub.2016.07.079)

10. Bayarri MJ, Muñoz-Cueto JA, López-Olmeda JF, Vera LM, Rol De Lama MA, Madrid JA, Sánchez-Vázquez FJ. 2004 Daily locomotor activity and melatonin rhythms in Senegal sole (Solea senegalensis). Physiol. Behav. 81, 577–583. (doi:10.1016/j.physbeh.2004.02.001)

11. Oliveira C, Garcia EM, López-Olmeda JF, Sánchez-Vázquez FJ. 2009 Daily and circadian melatonin release in vitro by the pineal organ of two nocturnal teleost species: Senegal sole (Solea senegalensis) and tench (Tinca tinca). Comp. Biochem. Physiol. - A Mol. Integr. Physiol. 153, 297–302. (doi:10.1016/j.cbpa.2009.03.001)

12. Huber R, Van staaden MJ, Kaufman LS, Liem KF. 1997 Microhabitat use, trophic patterns, and the evolution of brain structure in african cichlids. Brain. Behav. Evol. 50, 167–182. (doi:10.1159/000113330)

13. Albertson RC. 2008 Morphological Divergence Predicts Habitat Partitioning in a Lake Malawi Cichlid Species Complex. Copeia 2008, 689–698. (doi:10.1643/cg-07-217)

14. Parnell NF, Todd Streelman J. 2011 The macroecology of rapid evolutionary radiation. Proc. R. Soc. B Biol. Sci. 278, 2486–2494. (doi:10.1098/rspb.2010.1950)

15. Malinsky M et al. 2015 Genomic islands of speciation separate cichlid ecomorphs in an East African crater lake. Science (80-.). 350, 1493–1498. (doi:10.1126/science.aac9927)

16. Terai Y et al. 2017 Visual adaptation in Lake Victoria cichlid fishes: Depth-related variation of color and scotopic opsins in species from sand/mud bottoms. BMC Evol. Biol. 17. (doi:10.1186/s12862-017-1040-x)

17. Karvonen A, Wagner CE, Selz OM, Seehausen O. 2018 Divergent parasite infections in sympatric cichlid species in Lake Victoria. J. Evol. Biol. 31, 1313–1329. (doi:10.1111/jeb.13304)

18. Fryer G, Iles TD. 1972 cichlid fishes of the great lakes of Africa.

19. Arnegard ME, Carlson BA. 2005 Electric organ discharge patterns during group hunting by a mormyrid fish. Proc. R. Soc. B Biol. Sci. 272, 1305–1314. (doi:10.1098/rspb.2005.3101)

20. Siegel JM. 2009 Sleep viewed as a state of adaptive inactivity. Nat. Rev. Neurosci. 10, 747–753. (doi:10.1038/nrn2697)

21. Reebs SG, Colgan PW. 1991 Nocturnal care of eggs and circadian rhythms of fanning activity in two normally diurnal cichlid fishes, Cichlasoma nigrofasciatum and Herotilapia multispinosa. Anim. Behav. 41, 303–311. (doi:10.1016/S0003-3472(05)80482-8)

22. Reebs SG, Colgan PW. 1992 Proximal cues for nocturnal egg care in convict cichlids, Cichlasoma nigrofasciatum. Anim. Behav. 43, 209–214. (doi:10.1016/S0003-3472(05)80216-7)

23. Schwalbe MAB, Bassett DK, Webb JF. 2012 Feeding in the dark: Lateral-line-mediated prey detection in the peacock cichlid Aulonocara stuartgranti. J. Exp. Biol. 215, 2060–2071. (doi:10.1242/jeb.065920)

24. Jaggard JB, Lloyd E, Lopatto A, Duboue ER, Keene AC. 2019 Automated Measurements of Sleep and Locomotor Activity in Mexican Cavefish. J. Vis. Exp., e59198. (doi:10.3791/59198)

25. Yoshizawa M, Robinson BG, Duboué ER, Masek P, Jaggard JB, O’Quin KE, Borowsky RL, Jeffery WR, Keene AC. 2015 Distinct genetic architecture underlies the emergence of sleep loss and prey-seeking behavior in the Mexican cavefish. BMC Biol. 13. (doi:10.1186/s12915-015-0119-3)

26. Schneider CA, Rasband WS, Eliceiri KW. 2012 NIH Image to ImageJ: 25 years of image analysis. Nat. Methods. 9, 671–675. (doi:10.1038/nmeth.2089)

27. Howland HC, Merola S, Basarab JR. 2004 The allometry and scaling of the size of vertebrate eyes. Vision Res. 44, 2043–2065. (doi:10.1016/j.visres.2004.03.023)

28. R Core Team. 2017 R: A language and environment for statistical computing. R Foundation for Statistical. Vienna, Austria: R Foundation for Statistical Computing. See https://www.r-project.org/.

29. Maruyama A, Rusuwa B, Yuma M. 2010 Asymmetric interspecific territorial competition over food resources amongst Lake Malawi cichlid fishes. African Zool. 45, 24–31. (doi:10.1080/15627020.2010.11657251)

30. Ribbink A, Marsh B, Marsh A, Zoology AR-A, 1983 undefined. In press. A preliminary survey of the cichlid fishes of rocky habitats in Lake Malawi: results. ajol.info.

31. Konings A. In press. Malawi cichlids in their natural habitat. 3rd edn. Cichlid Press. El Paso, Texas

32. Willacker JJ, Von Hippel FA, Wilton PR, Walton KM. 2010 Classification of threespine stickleback along the benthic-limnetic axis. Biol. J. Linn. Soc. 101, 595–608. (doi:10.1111/j.1095-8312.2010.01531.x)

33. Archer S. 1999 Adaptive Mechanisms in the Ecology of Vision. Boston, MA: Kluwer Academic Publishers.

34. Motani R, Rothschild BM, Wahl W. 1999 Large eyeballs in diving ichthyosaurs. Nature 402, 747. (doi:10.1038/45435)

35. Hulsey CD, Mims MC, Streelman JT. 2007 Do constructional constraints influence cichlid craniofacial diversification? Proc. R. Soc. B Biol. Sci. 274, 1867–1875. (doi:10.1098/rspb.2007.0444)

36. Cooper WJ, Westneat MW. 2009 Form and function of damselfish skulls: Rapid and repeated evolution into a limited number of trophic niches. BMC Evol. Biol. 9, 24. (doi:10.1186/1471-2148-9-24)

37. Genner MJ, Seehausen O, Cleary DFR, Knight ME, Michel E, Turner GF. 2004 How does the taxonomic status of allopatric populations influence species richness within African cichlid fish assemblages? J. Biogeogr. 31, 93–102. (doi:10.1046/j.0305-0270.2003.00986.x)

38. Barlow G. 2008 The cichlid fishes: nature’s grand experiment in evolution. See https://books.google.com/books?hl=en&lr=&id=LME6bc4m6QcC&oi=fnd&pg=PR9&dq=barlow+cichlid&ots=i_LH7pD9ai&sig=6cMb8AC7RHPbdY3D4JInvqGvkyY.

39. Vera LM, Cairns L, Sánchez-Vázquez FJ, Migaud H. 2009 Circadian Rhythms of Locomotor Activity in the Nile Tilapia *Oreochromis niloticus*. Chronobiol. Int. 26, 666–681. (doi:10.1080/07420520902926017)

40. del Pozo A, Sánchez-Férez JA, Sánchez-Vázquez FJ. 2011 Circadian Rhythms of Self-feeding and Locomotor Activity in Zebrafish (*Danio Rerio*). Chronobiol. Int. 28, 39–47. (doi:10.3109/07420528.2010.530728)

41. Reebs SG. 1994 Nocturnal Mate Recognition and Nest guarding by Female Convict Cichlids (Pisces, Cichlidae: Cichlasoma mgrofasciatum). Wiley Online Libr. 96, 303–312. (doi:10.1111/j.1439-0310.1994.tb01018.x)

42. Reebs SG. 2002 Plasticity of diel and circadian activity rhythms in fishes. Rev. Fish Biol. Fish. 12, 349–371. (doi:10.1023/A:1025371804611)

43. Won YJ, Sivasundar A, Wang Y, Hey J. 2005 On the origin of Lake Malawi cichlid species: A population genetic analysis of divergence. Proc. Natl. Acad. Sci. U. S. A. 102, 6581–6586. (doi:10.1073/pnas.0502127102)

44. Conith MR, Hu Y, Conith AJ, Maginnis MA, Webb JF, Craig Albertson R. 2018 Genetic and developmental origins of a unique foraging adaptation in a Lake Malawi cichlid genus. Proc. Natl. Acad. Sci. U. S. A. 115, 7063–7068. (doi:10.1073/pnas.1719798115)

45. Conith MR, Conith AJ, Albertson RC. 2019 Evolution of a soft-tissue foraging adaptation in African cichlids: Roles for novelty, convergence, and constraint. Evolution (N. Y). 73, 2072–2084. (doi:10.1111/evo.13824)

46. Albertson RC, Pauers MJ. 2019 Morphological disparity in ecologically diverse versus constrained lineages of Lake Malaŵi rock-dwelling cichlids. Hydrobiologia 832, 153–174. (doi:10.1007/s10750-018-3829-z)

47. Keene AC, Duboue ER. 2018 The origins and evolution of sleep. J. Exp. Biol. 221. (doi:10.1242/jeb.159533)

48. Prober DA, Rihel J, Onah AA, Sung RJ, Schier AF. 2006 Hypocretin/orexin overexpression induces an insomnia-like phenotype in zebrafish. J. Neurosci. 26, 13400–13410. (doi:10.1523/JNEUROSCI.4332-06.2006)

49. Duboué ER, Keene AC, Borowsky RL. 2011 Evolutionary Convergence on Sleep Loss in Cavefish Populations. Curr. Biol. 21, 671–676. (doi:10.1016/J.CUB.2011.03.020)

50. Aspiras AC, Rohner N, Martineau B, Borowsky RL, Tabin CJ. 2015 Melanocortin 4 receptor mutations contribute to the adaptation of cavefish to nutrient-poor conditions. Proc. Natl. Acad. Sci. U. S. A. 112, 9668–73. (doi:10.1073/pnas.1510802112)

