## Supplemental Figure 1 for "Diversity in rest-activity patterns among Lake Malawi cichlid fishes suggests novel axis of habitat partitioning"

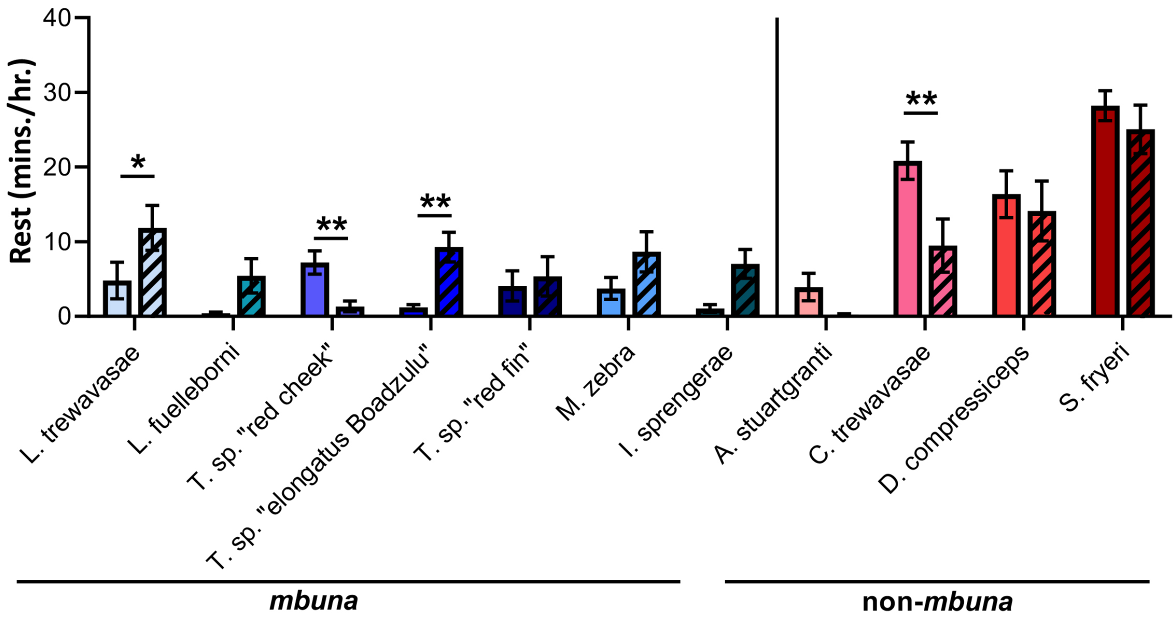


**Fig. S1.** Cichlid species demonstrate interspecies variation in activity/rest cycles throughout the day.
